# The BioLuminescent-OptoGenetic *in vivo* Response to Coelenterazine is Proportional, Sensitive and Specific in Neocortex

**DOI:** 10.1101/709931

**Authors:** Manuel Gomez-Ramirez, Alexander I. More, Nina G. Friedman, Ute Hochgeschwender, Christopher I. Moore

**Author notes:** Correspondence: Christopher I. Moore, Ph.D, Department of Neuroscience, Brown University, 185 Meeting Street. Providence, RI 02912.

## Abstract

BioLuminescent (BL) light production can modulate neural activity and behavior through coexpressed OptoGenetic (OG) elements, an approach termed ‘BL-OG’. Yet, the relationship between BL-OG effects and bioluminescent photon emission has not been characterized *in vivo*. Further, the degree to which BL-OG effects strictly depend on optogenetic mechanisms driven by bioluminescent photons is unknown. Crucial to every neuromodulation method is whether the activator shows a dynamic concentration range driving robust, selective, and non-toxic effects. We systematically tested the effects of four key components of the BL-OG mechanism (luciferin, oxidized luciferin, luciferin vehicle, and bioluminescence), and compared these against effects induced by the Luminopsin-3 (LMO3) BL-OG molecule, a fusion of slow burn Gaussia luciferase (sbGLuc) and Volvox ChannelRhodopsin-1 (VChR1). We performed combined bioluminescence imaging and electrophysiological recordings while injecting specific doses of Coelenterazine (substrate for sbGluc), Coelenteramide (CTM, the oxidized product of CTZ), or CTZ vehicle. CTZ robustly drove activity in mice expressing LMO3, with photon production proportional to firing rate. In contrast, low and moderate doses of CTZ, CTM, or vehicle did not modulate activity in mice that did not express LMO3. We also failed to find bioluminescence effects on neural activity in mice expressing an optogenetically non-sensitive LMO3 variant. We observed weak responses to the highest dose of CTZ in control mice, but these effects were significantly smaller than those observed in the LMO3 group. These results show that in neocortex in vivo, there is a large CTZ range wherein BL-OG effects are specific to its active chemogenetic mechanism.

## INTRODUCTION

Novel genetic- and optical-based methods that target specific cell types provide a powerful strategy for probing neural mechanisms that underlie perception and action (Boyden et al., 2005; Zhang et al., 2006; Cardin et al., 2009; Nichols and Roth, 2009; Knopfel et al., 2010; Fenno et al., 2011; Tung et al., 2015; Zhu et al., 2015; Berglund et al., 2016a; Roth, 2016; Kim et al., 2017). Of such approaches, optogenetic and chemogenetic strategies are the most widely employed. The former provides high spatio-temporal precision, while the latter provides broad coverage and minimally-invasive delivery of the activating driver.

BioLuminescent OptoGenetics (‘BL-OG’) is an emerging dual strategy that provides both optogenetic and chemogenetic capabilities. In this approach, binding of an oxidative enzyme (luciferase) to a small light-emitting molecule (luciferin) drives bioluminescence that, in turn, regulates a neighboring opsin (Berglund et al., 2013; Birkner et al., 2014; Tung et al., 2015; Berglund et al., 2016b; Berglund et al., 2016a; Park et al., 2017; Prakash et al., 2018; Tung et al., 2018; Zenchak et al., 2018; Berglund et al., 2019). To ensure proximity of the bioluminescent reaction to the recipient opsin, the *luminopsin* (LMO) construct was invented, in which a luciferase is linked to the optogenetic element by a short 15 amino acids linker (Berglund et al., 2016a). Here, we use LMO3, a molecule that tethers the slow-burn Gaussia luciferase (sbGLuc) to Volvox Channelrhodopsin-1 (VChR1; Figure 1A), and uses the substrate coelenterazine (CTZ) to generate bioluminescence. This single molecule permits both chemogenetic regulation by peripheral injection of a luciferin (Berglund et al., 2013; Birkner et al., 2014; Tung et al., 2015; Berglund et al., 2016a), and local optogenetic regulation by external light application or direct intracortical injection of the luciferin (Tung et al., 2015). The BL-OG strategy has several additional distinctive benefits, including the robust biocompatibility of its components, and photon production that confirms the activator reached its target (e.g., by imaging while administering the drug). Further, while existing BL-OG implementations have employed luciferases that bind the luciferin Coelenterazine (CTZ), multiple distinct and non-interacting classes of luciferin are potentially viable (Haddock et al., 2010), providing multiple independent pathways for distinct forms of simultaneous modulation.

**Figure 1.**
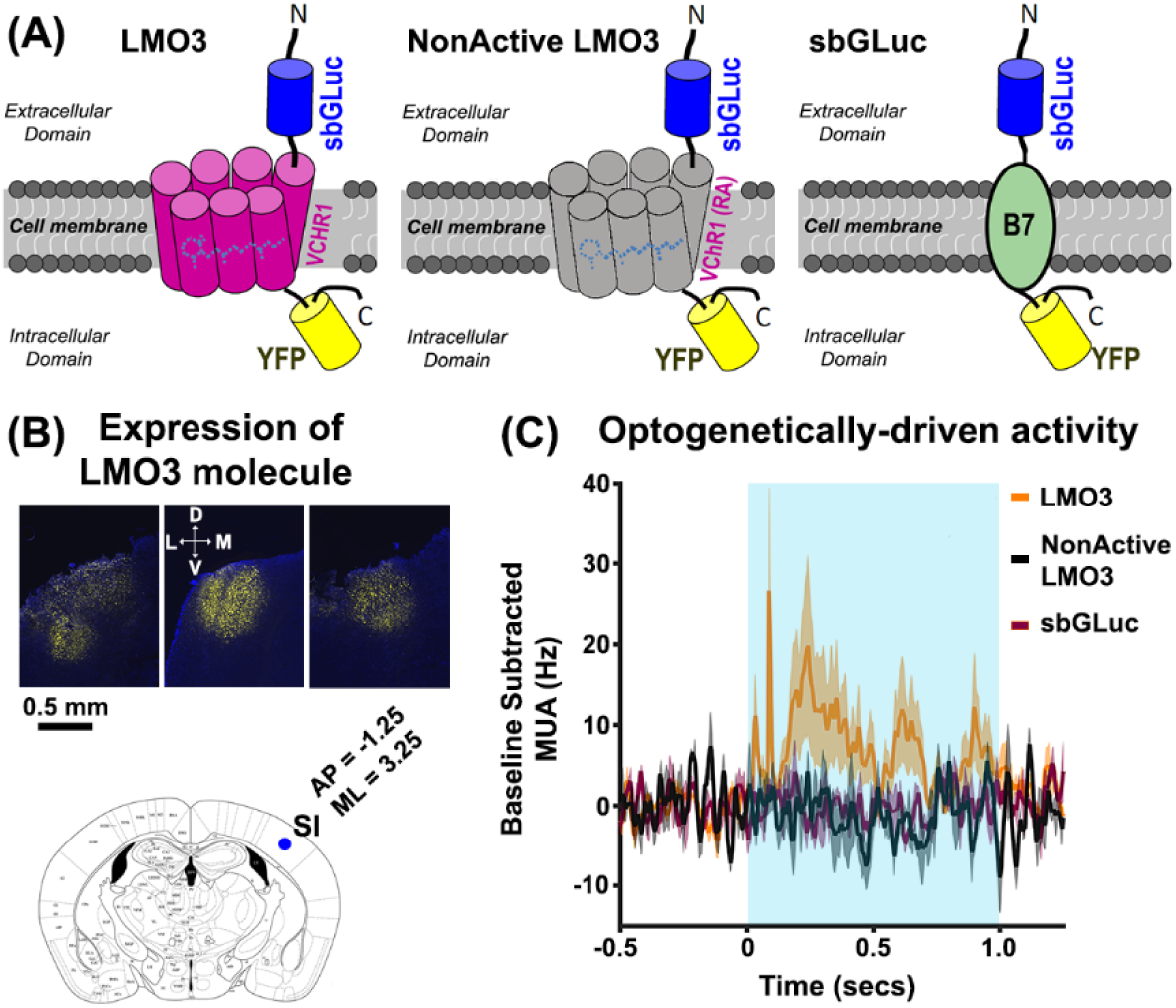
**(A)** Schematic illustration of the genetically-engineered molecules: LMO3 (left), NonActive LMO3 (center), and sbGLuc (right). In the LMO3 molecule, the luciferase slow-burn Gaussia (sb-GLuc) is tethered to the opsin Volvox Channelrhodopsin1 (VChR1) and the enhanced yellow fluorescent protein (eYFP). The NonActive LMO3 is structurally the same as the LMO3 molecule, except that the VChR1 has an arginine-alanine point mutation that renders the opsin nonfunctional, indicated by the gray color of the VChR1. The sbGLuc is linked to the membrane using the B7 transmembrane protein with the luciferase located outside the cell. **(B)** Representative histological images showing eYFP expression in the left SI of three animals (upper panels). The lower panel shows a brain slice of a mouse atlas at location A/P = -1.25, M/L = 3.25 relative to Bregma. The blue circle indicates the depth of the injection site (∼500μm). We injected in three locations relative to the blue circle: Location 1 (A/P = -0.75 M/L = 2.75), Location 2 (A/P = 0.5, M/L = 0), Location 3 (A/P = -1.75 M/L = 2.75). All injections were made the same depth (500 μm) (mouse brain atlas was adapted from http://labs.gaidi.ca/mouse-brain-atlas/). **(C)** Spiking activity in the LMO3 (orange trace), NonActive LMO3 (black trace), and sbGLuc (violet trace) groups in response to LED stimulation. The data show increases in spiking activity for the LMO3 group only, indicating that only this cohort expressed a functional optogenetic element.

Activators (e.g., chemicals or light) used in all neuromodulation methods have a regime of non-specificity. The most widely-employed chemogenetic method, Designer Receptors Exclusively Activated by Designer Drugs (DREADDs), is driven by clozapine n-oxide (CNO) (Urban and Roth, 2015). MacLaren and colleagues (2016) showed that CNO has adverse behavioral effects in naïve rats, possibly because CNO is converted into Clozapine (an antipsychotic) through endogenous body mechanisms (Gomez et al., 2017). Light stimulation can also impact local cellular processes by a variety of intrinsic mechanisms, including photovoltaic/photoelectric effects (Kozai and Vazquez, 2015). A crucial challenge for neuromodulation methods is to determine a dose (or intensity) range of the activator that evokes a proportional modulation, so that a desired effect is consistently achieved throughout repeated trials.

Here, we systematically tested the relationship between BL-OG neural modulation and photon production, and the possibility that non-specific components of its bioluminescent reaction modulate neuronal spiking. To this end, we expressed LMO3 in mice, a BL-OG construct previously demonstrated to robustly activate neurons (Berglund et al., 2016b; Berglund et al., 2016a; Prakash et al., 2018), and drive behavioral responses when activated by CTZ (Berglund et al., 2016a; Zenchak et al., 2018). We tested whether the neural response in LMO3-expressing mice was proportional to CTZ dose and photon production. The BL-OG mechanism requires catalysis of CTZ, which generates its oxidized derivative, Coelenteramide (CTM). As such, we also tested the effects of CTZ, CTM, or vehicle control (the compound that renders CTZ water soluble) on neural spiking in naïve mice. We also assayed the effects of bioluminescence itself on neural activity by injecting CTZ in mice expressing sbGLuc without a functional opsin or the sbGLuc alone. BL-OG effects in mice expressing LMO3 were highly sensitive, showing proportional increases in neural firing and photon production in response to increasing doses of CTZ. Systematic responses were not observed in any control condition. We only observed weak, but significant, responses to the highest CTZ dose in naïve mice and mice not expressing a functional opsin. These results show a selective and robust regime for driving neocortical activation with the BL-OG method.

## METHODS

### Animals

Thirty-six mice (16 females, 20 males, C57BL/6NHsd from Jackson Labs, 10-26 weeks) were utilized in the experiments. Mice were housed in a vivarium with reverse light-dark cycle (12 hours each phase) and given *ad libidum* access to water and food. All procedures were conducted in accordance with the guidelines of the National Institute of Health and with approval of the Animal Care and Use Committee of Brown University.

### BL-OG Construct and Viral Injections

Figure 1A shows schematics of each genetically-engineered molecule used in this paper, and Table 1 describes the different molecules and chemical agents injected in each cohort. The LMO3 molecule (sbGLuc-VChR1-EYFP) (Berglund et al., 2016b) consists of human codon optimized Gaussia luciferase carrying two methionine-to-leucine mutations (M43L/M110L; sbGLuc) (Welsh et al., 2009) fused to the extracellular N-terminus of Volvox Channelrhodopsin 1 (VChR1-EYFP) (Zhang et al., 2008). For the NonActive LMO3 (R115A) construct, a non-functional arginine-to-alanine point mutation described for ChR2 (R120A) (Kato et al., 2012) was introduced into the VChR1 sequence. All molecules were packaged in an AAV2/9 vector with the human synapsin 1 promoter. For the GLuc-only construct, the sequence coding for VChR1 was replaced by the sequence for the B7 transmembrane region from the mouse CD80 antigen (Chou et al., 1999), generating sbGLuc-B7-EYFP. The luciferase is tethered to the cell membrane extracellularly without the optogenetic element.

**Table 1:** Description of the molecule each cohort expressed and the chemical agents injected.

| Table 1 |  |  |
| --- | --- | --- |
| Cohort | Chemical injected | Molecule expression |
| LMO3 | CTZ | LMO3 |
| CTM | CTM | None |
| Vehicle | vehicle | None |
| GLuc | CTZ | NonActive-LMO3<br>sbGLuc-B7 |
| CTZ | CTZ | none |

We injected 450nl of the virus across three locations in left primary somatosensory (SI) cortex (150nl per site). Injections were made through a burr hole placed 500μm M/L and 500μm A/P relative to Bregma in SI (−1.25 A/P and 3.25 M/L relative to Bregma). Specifically, injection 1 was made at A/P = -1.25 M/L = 3.75, injection 2 was made at A/P = -0.75 M/L = 2.75. and injection 3 was made A/P = -1.75 M/L = 2.75. Figure 1B shows examples of viral expression data, and an atlas illustration of the injections. The virus was drawn into a glass micropipette attached to a Quintessential Stereotaxic Injector (QSI, Stoelting) that was lowered 500μm below the cortical surface. Viral constructs were infused at a rate of 7.5nl/min, and pipettes were held in place for 5 minutes following infusions before retracting from the brain. The scalp incision was manually sutured or glued using Gluture (Abbott Laboratories). Dexamethasone was given intraperitoneally (0.1 mg/kg) to reduce brain inflammation, and Slow Release Buprenex was given subcutaneously to aid post-surgery comfort (0.1 mg/kg). The median time between viral transduction surgery and experiments was 28 days (min = 26 days, max time = 74 days). There was no systematic relationship between expression time and bioluminescence (p > 0.05; r = 0.007) or MUA (p > 0.05; r = 0.021).

### Luciferin injections

Water soluble CTZ (Nanolight Technology, Catalog #3031 Coelenterazine-SOL *in vivo)* was diluted in sterile water (1mg/ml) to yield a concentration of 2.36mM. CTZ injections were done directly in cortex. In a preliminary subset of LMO3-expressing mice, we also assayed BL-OG effects by injecting CTZ via intraperitoneal, intravenous, and intraventricular routes (data not shown). Water soluble CTM was purchased from Nanolight Technology, and diluted in sterile water (1 mg/ml) to yield a concentration of 2.43mM. The vehicle was also purchased from Nanolight Technology, and diluted in sterile water (1mg/ml). Local cortical injections of CTZ, CTM, and vehicle spanned 0.2μl, 0.4 μl, and 1μl, at an injection rate of 1.25μL/min. All cortical injections were delivered using a Hamilton Nanosyringe with a 34-gauge needle that was attached to a motorized injector (Stoelting Quintessential Stereotaxic Injector, QSI).

### Imaging and Electrophysiology Recordings

Bioluminescence was measured using an electron multiplier charge-coupled-device (EMCCD) camera (Ixon 888, Andor) attached to a Navitar Zoom 6000 lens system (Navitar, 0.5x lens). The camera’s sensor was 1024 × 1024 pixels, with a 13.4μm x 13.4μm pixel size. Images were collected in a custom-made light-tight chamber with an exposure time of 10 seconds, and the EM gain set to 30. Imaging data were recorded using the Solis image acquisition and analysis software (Solis 4.29, Andor). A TTL pulse was used to synchronize the onset of imaging and electrophysiological data in each experimental block.

Neurophysiological activity was recorded using a 32 channel laminar electrode (Optoelectrode, A1×32-Poly2-5mm-50s-177-OA32LP, Neuronexus). The laminar electrode was composed of two columns of 16 channels with each contact spaced 50μm apart. The electrode had an optical fiber (105μm diameter) attached to the recording side of the electrode shank that terminated 200μm above the most superficial electrode. The electrode was lowered 900μm past the cortical surface. Neural activity was acquired using the open source OpenEphys system (http://www.open-ephys.org/). Neural data were recorded with a sampling rate of 20KHz, and referenced to a supra-dural electrode chronically implanted over right occipital cortex.

### Optogenetic Stimulation

Optogenetic stimulation was achieved by delivering 40Hz light pulses of 470nm wavelength. Each pulse intensity was 0.5 mW/cm2, 0.07 seconds in length, and with the probability drawn from a Poisson distribution (rate = 40Hz). The entire 40Hz pulse train lasted one second. Light was controlled using a Mightex LED driver (SLA-1200-2) that delivered power to a Thorlabs fiber-coupled LED. A patch cable from the fiber-coupled LED was attached to a bare ferrule-coupled connector in the optoelectrode.

### Sequence of Events

Experiments were performed under isoflurane delivered at ∼1% (range 0.75% to 2%). Approximately 1 hour prior to recording, a craniotomy that covered virally-transduced sites was performed, and animals were transferred from the surgical suite to the experimental chamber. The order of CTZ cortical injection doses was randomized across animals. Each session began with 10 seconds of baseline recordings. Sensory vibrissal stimuli were applied before and after CTZ injection (3-5 seconds inter-stimulus interval; ISI). Note however, that analyses were conducted on activity during the pre-stimulus because our goal was to assay BL-OG effects uncontaminated by sensory-driven activity. Data associated with tactile stimulation are not reported as receptive fields were inconsistent or absent in a subset of recordings.

In a subset of animals expressing LMO3 (N = 7), we performed optogenetic studies to assay whether opsins were driven using standard optical methods. We performed similar experiments in animals expressing the sbGLuc-only and the NonActive-LMO3 molecule. Figure 1C shows mean activity in response to LED activity in the 470nm range for animals expressing LMO3 (orange trace), sbGLuc-only (violet trace), and Non-Active LMO3 (black trace). The data show activation in LMO3 mice only, indicating that the VChR1 opsin was functional in this cohort.

### Histology

After each experiment, mice were euthanized with isoflurane and perfused transcardially with 4% paraformaldehyde (PFA). The brain was removed, and kept in PFA at 4°C for 36 hours after perfusion. The brain was cryoprotected in 30% sucrose for another 36 hours prior to tissue slicing. Mice brains were sectioned in 50μm slices on a cryostat (Leica CM30505), and mounted on glass slides for imaging on an inverted fluorescent microscope (Zeiss Axiovert 200M; 10x EC Plan-NeoFluar objective). Regions of viral expression were then compared to a brain atlas (Allen Mouse Brain Atlas) to confirm the correct location of the SI viral expression (see Figure 1B).

### Analyses

Images were converted to 16-bit tiff files and analyzed using custom-based scripts in MATLAB. Pixel values higher than three standard deviations from the mean present for only a single frame were deleted, and images were smoothed using a 2×2 pixel filter. For each pixel, bioluminescence was expressed as a relative measure by subtracting the averaged activity between the initial frame and the frame prior to CTZ injection from all recorded frames. Statistical significance of bioluminescent activity was assayed using non-parametric bootstrapping statistics. For each pixel, we computed the mean of 25 randomly extracted frames prior to the CTZ injection, and built a surrogate distribution by repeating this procedure 5000 times. Bioluminescent activity of each pixel after CTZ injection was compared against the surrogate distribution. Significant activity was defined as data values greater than 95% of the surrogate values for at least 3 consecutive frames.

Electrophysiological data were analyzed using custom scripts in MATLAB. Data were downsampled (n = 2) and notch filtered at 60Hz to reduce ambient noise. Proxies of multi-unit activity (MUA) were derived by filtering the raw data with a high-pass filter (500 and 2500Hz), and rectifying the filtered data. Outliers were removed by deleting data points nine standard deviations greater than the average power value. A value of one (i.e., a spike) was assigned to data points that were four standard deviations greater than the mean. All other points were assigned a zero value. We convolved MUA activity with an asymmetric Gaussian filter (30ms duration, with a rising slope = 19.91mV/s) to derive instantaneous MUA (Gomez-Ramirez et al., 2014). Individual observations used in the statistics comprised averaged activity between - 1500ms and -500ms prior to the onset of the tactile stimulus. Statistical effects were assayed using non-parametric bootstrapping statistics, with significant activity defined as values greater than 95% of the surrogate values. The data that support the findings of this study are available from the corresponding author upon reasonable request.

## RESULTS

### Bioluminescent photon generation is proportional to neural firing in LMO3-expressing mice

We performed simultaneous electrophysiology and high-sensitivity imaging in primary somatosensory neocortex (SI) in mice transduced with LMO3 (N = 10). Figure 2A shows an example of enhanced bioluminescence following CTZ injection. Figure 2B shows the bioluminescence and MUA time courses for the ROI depicted in Figure 2A (*dashed box*) near the electrode (*white circle*) and injection cannula (*red circle*).

**Figure 2.**
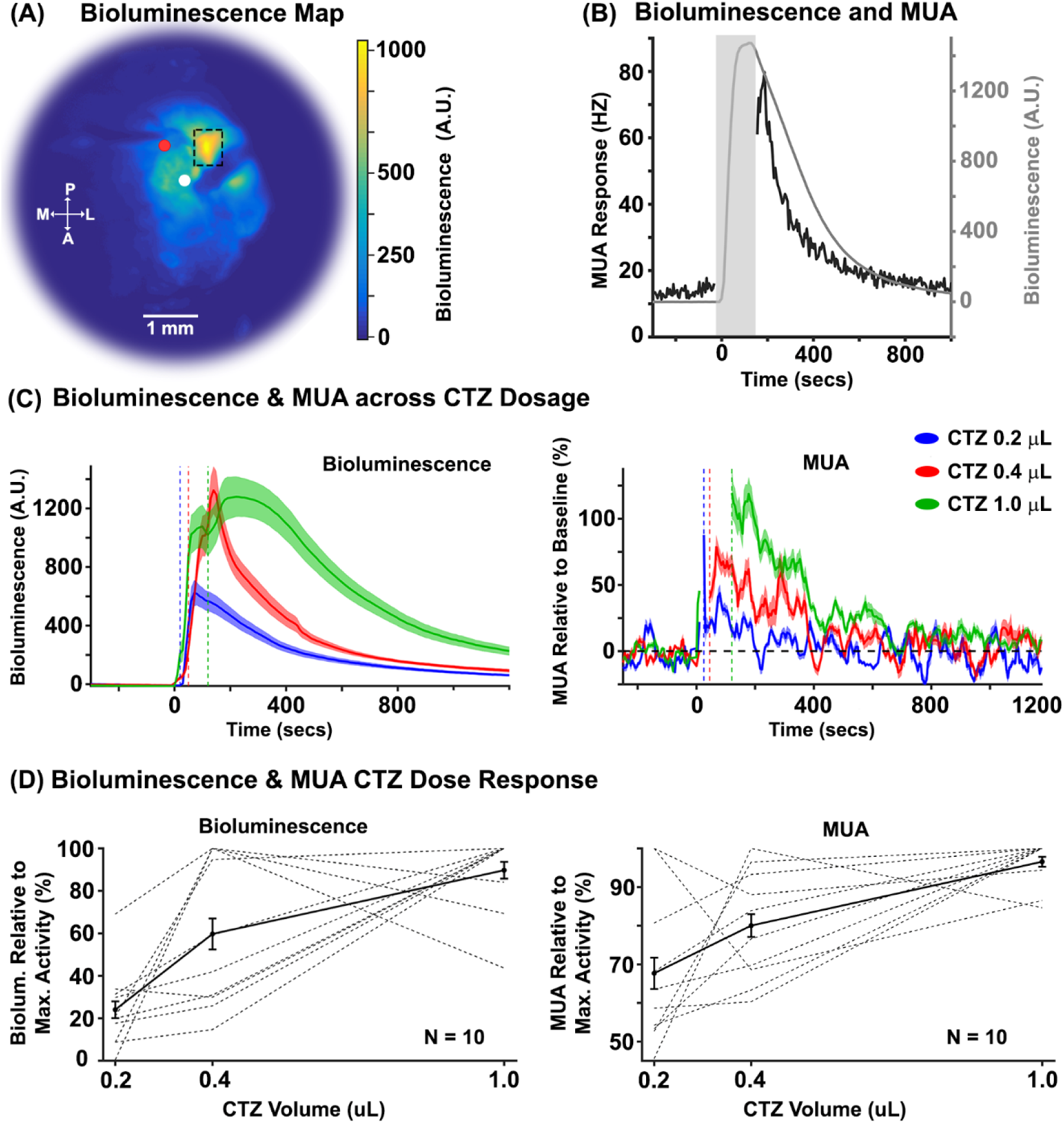
BL-OG effects as a function of CTZ dose. **(A)** Representative example of a normalized bioluminescence color map in response to a 1.0 μL injection of CTZ directly in cortex. A = anterior, P = posterior, M = medial, L = lateral. The white and red circles indicate the surface locations of the electrode and injector, respectively. The electrode and syringe were inserted at angles between ∼20° and 45°, and to different depths relative to dura (electrode ∼900μm, and syringe ∼500μm). **(B)** Bioluminescence (gray trace) and MUA (black trace) time courses within the ROI in Figure 2A (dashed box). The gray rectangle indicates the time of CTZ onset/offset for the 1.0 μL CTZ dose condition with a +/-30 seconds window added. The discontinuity in the MUA response stemmed from a mechanical and electrical artifact created by the QSI injector. **(C)** Left and right panels show normalized bioluminescence and MUA as a function of CTZ dose for direct cortical injection conditions, respectively (N = 10). The dotted vertical lines indicate the offset of the injection for each CTZ dose condition. **(D)** Left and right panels show normalized bioluminescence and MUA for each animal, relative to the CTZ dose that evoked the max response, respectively (N = 10).

To systematically compare the sensitivity and proportionality of the optical and neurophysiological response, we injected three CTZ doses (0.2, 0.4, and 1.0 μL) (Figure 2C). All dose levels of CTZ significantly increased bioluminescence relative to baseline at the mouse group level (p <0.0001, mean 0.2 μL = 419.75, mean 0.4 μL = 715.15, mean 1.0 μL = 1233.14;), as well as in each mouse (p < 0.0001), with the exception of two mice at the lowest dose (p = 0.15 and p = 0.101). The CTZ injection also increased MUA at the group level. Specifically, we found significant increases in MUA for the 0.4 μL (mean = 41.39%; p = 0.039), and 1.0 μL (mean = 71.22%; p = 0.01) CTZ doses. We observed a trend towards significance for the 0.2 μL condition (mean = 22.48%; p = 0.09). At the individual mouse level, we observed significant modulations in MUA in six mice for 0.2 μL CTZ (p < 0.05), eight mice for 0.4 μL CTZ (p < 0.05), and nine mice for 1.0 μL CTZ (p < 0.05). The left and right panels of Figure 2D show bioluminescence (left panel) and MUA (right panel) responses normalized to the highest CTZ response of each animal, respectively. Linear regression analyses revealed a systematic relationship between bioluminescence and MUA (p < 0.001, R^2^ = 0.36; see Figure 3; gray regression line).

**Figure 3.**
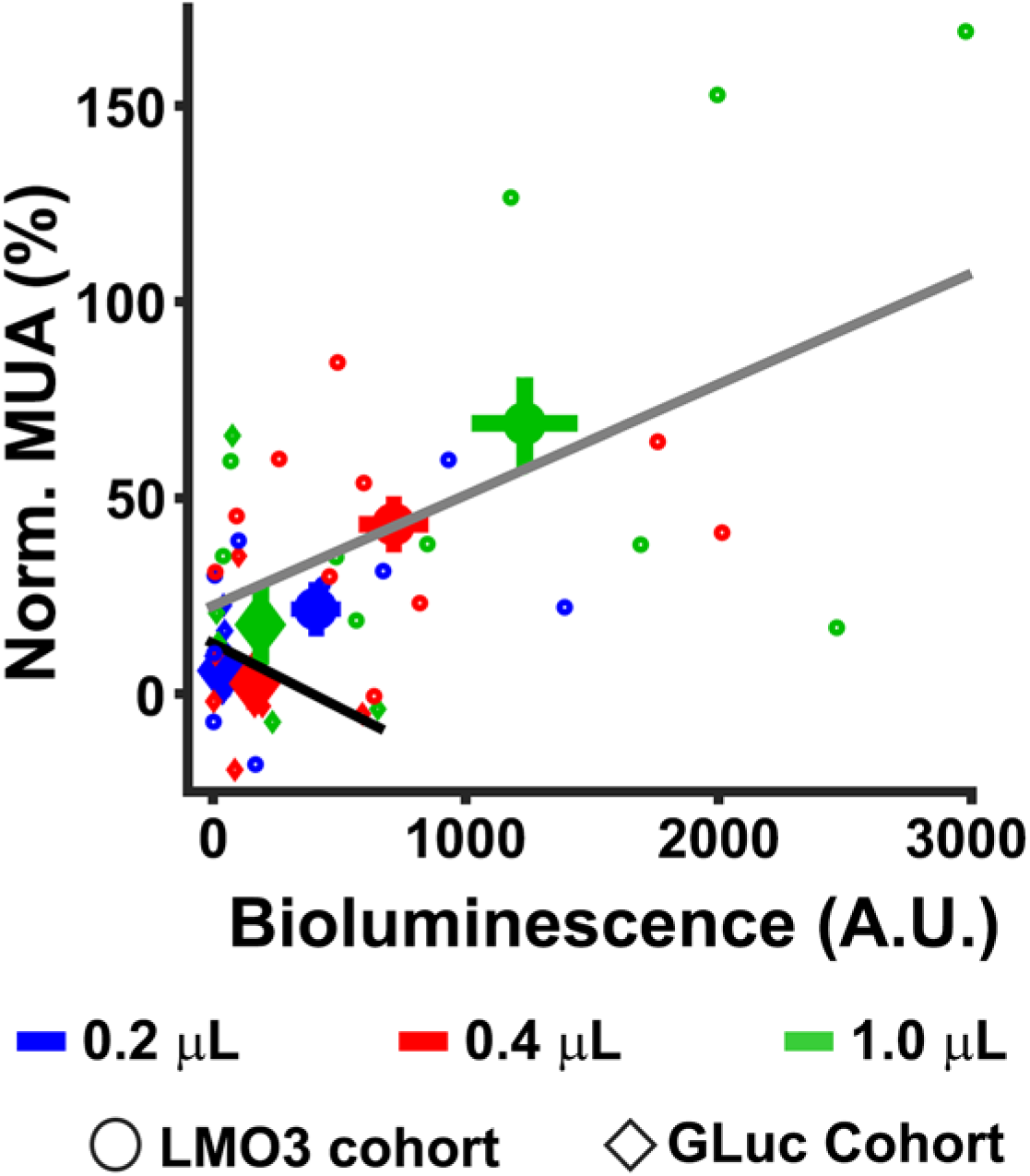
MUA as a function of bioluminescence. The relationship is shown between averaged bioluminescence and MUA for each CTZ dose condition in the LMO3 (circle symbol) and GLuc (diamond symbol) cohorts for the time period (0 – 400 seconds after injection of CTZ). The larger symbols indicate the mean activity for each CTZ dose condition.

We failed to see significant modulations in MUA at the mouse group level during later time periods (400 to 800 seconds) for any CTZ dose condition (0.2 μL = 0.04%, p > 0.05; 0.4 μL = 7.90%, p > 0.05; 1.0 μL = 22.82%, p > 0.05). However, at the individual mouse level, we observed significant MUA increases in two mice for 0.2 μL (p < 0.05), two mice for 0.4 μL CTZ (p < 0.05), and four mice for 1.0 μL CTZ (p < 0.05). Importantly, we observed significant increases in bioluminescence during the latter periods (400 to 800secs) for all CTZ doses (0.2 μL = 154.78, p < .0001; 0.4 μL = 232.24, p < .0001; 1.0 μL = 671.99, p < .0001) at the group level. At the individual mouse level, we observed significant bioluminescence in nine mice for 0.2 μL CTZ (p < 0.05), nine mice for 0.4 μL CTZ (p < 0.05), and ten mice for 1.0 μL CTZ (p < 0.05). This discrepancy between bioluminescence and MUA effects could either reflect weak bioluminescent light that is unable to drive detectable neural activity, adaptation of the neural response, and/or adaptation of optogenetic sensitivity to sustained bioluminescent production.

### Elements of the BL-OG reaction create minimal off target effects in vivo

We assayed whether CTZ, CTM, or vehicle drive spiking activity through non-BL-OG related mechanisms, by injecting these chemicals directly in SI of non-expressing mice (Figure 4A *upper three panels*). For the CTZ cohort (N = 8), we did not observe significant changes in MUA to the .2 μL (2.89%, p > 0.05) or 0.4 μL CTZ doses (2.26% p > 0.05). However, we found modest, but significant, MUA increases when injecting the largest dose of CTZ (1.0 μL = 12.79%, p = 0.045). For the CTM cohort (N = 8), no significant modulations for any dose conditions were observed (0.2 μL = -2.89%, p > 0.05; 0.4 μL = 0.63%, p > 0.05; 1.0 μL = 5.71%, p = 0.09). We also failed to find significant MUA changes in the vehicle cohort for any dose (0.2 μL = 0.4%, p > 0.05; 0.4 μL = 2.59%, p > 0.05; 1.0 μL = 2.77%, p > 0.05). As expected, we also failed to observe bioluminescent responses to any dose condition for the CTZ, CTM, and vehicle cohorts (p > 0.05).

**Figure 4.**
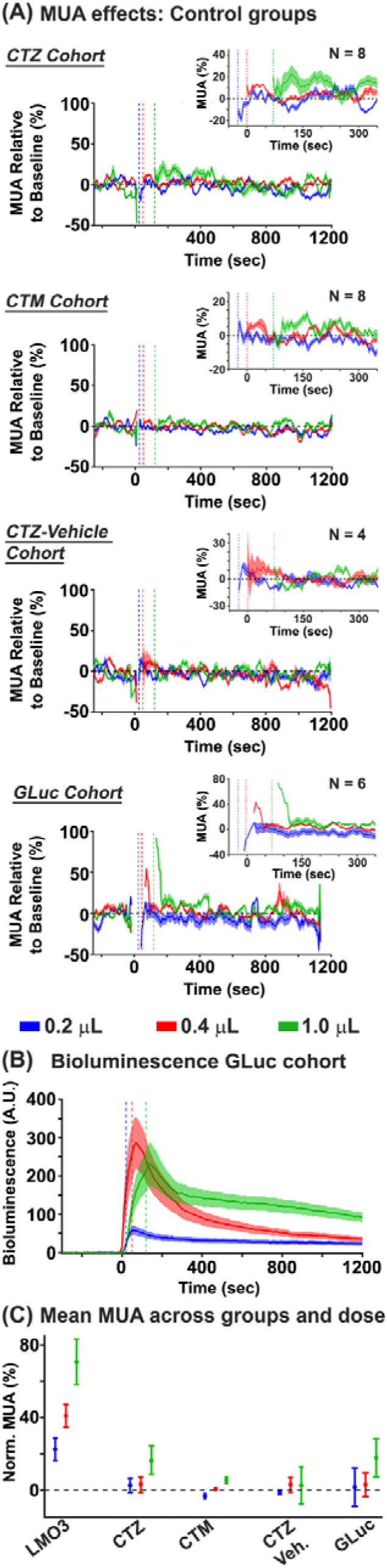
MUA and bioluminescence in control groups. **(A)** BL-OG effects on MUA are shown for the control groups. The top graph shows MUA effects in the CTZ naïve group. The data revealed significant increases in the 1.0μL condition. The second and third row graph shows MUA data in the CTM and vehicle naïve groups, respectively. The data did not reveal MUA changes for any dose condition in either group. The bottom row shows MUA changes for the GLuc cohort. Similar to the CTZ naïve group, we observed modest increases in MUA for the largest CTZ dose. The inset in all graphs shows an enlarged view of activity between -50 and 350secs after chemical injection. **(B)** The bioluminescent response across CTZ doses for the GLuc cohort is shown. Systematic increases in bioluminescence were observed with increases in CTZ dose. **(C)** MUA activity across all cohorts and chemical doses. The strongest MUA modulations were observed in the LMO3 group, with small changes for the largest CTZ dose for the CTZ naïve and GLuc groups.

To test for effects of bioluminescence on spiking activity, we injected a molecule that expressed sbGLuc or an LMO3 molecule with a mutation in the opsin that rendered the VChR1 nonfunctional in different animals. MUA responses in these conditions were similar, and not statistically different from each other (sbGLuc: 0.2 μL = 6.73%, 0.4 μL = -0.24%, 1.0 μL = 17.85%; NonActiveLMO3: 0.2 μL = -3.74%, 0.4 μL = 5.97%, 1.0 μL = 17.62%), thus we combined the two datasets, a cohort we termed ‘GLuc’ (Figure 4A *lower panels*, and Figure 4B). Injections of CTZ yielded no MUA changes in the 0.2 μL (1.49%, p > 0.05) or 0.4 μL CTZ dose (2.87%, p > 0.05). However similar to the CTZ cohort, we found modest significant increases in the 1.0 μL CTZ condition (17.74%, p < 0.01). We observed proportional modulations in bioluminescence in response to CTZ (0.2 μL = 38.52 p < 0.05; 0.4 μL = 165.48 p < 0.05; 1.0 μL = 188.58 p < 0.05) (see Figure 4B), but, importantly, failed to see a relationship between MUA and bioluminescent photon production (R^2^ = 0.09; p > 0.05, see Figure 3, black regression line). We compared MUA effects between the CTZ and GLuc cohorts to assay whether the slight increase in MUA for the GLuc vs. CTZ cohort (17.74% vs. 16.28%) was due to the bioluminescent reaction or just unaccounted noise. The data revealed no significant differences between the two groups (p = 0.61), indicating that MUA modulations in the GLuc group are likely driven by the CTZ chemical itself. A summary of the mean MUA effects for each chemical dose condition and cohort tested is shown in Figure 4C.

### Temporal properties of the BL-OG effect

We investigated the duration of the BL-OG effect by determining the continuity of significant MUA increases for all CTZ doses in the LMO3 group. We classified a statistically significant time point when two consecutive time points had an increase in MUA greater than 95% of points as compared to a surrogate distribution built from the CTZ pre-injection data (Figure 5). The data revealed continuous BL-OG effects for the largest CTZ dose that commenced at the earliest time point analyzed after CTZ onset, and lasted for ∼160 seconds (see inset Figure 5 green trace). Similarly, continuous BL-OG effects in the 0.4 μL started right after CTZ injection, but only lasted ∼40 seconds (red trace in Figure 5 inset). However, there was a long period (∼120 seconds) where MUA BL-OG effects in the 0.4 μL condition trended towards significance (p < 0.1). Continuous BL-OG effects in the 0.2 μL CTZ condition were not observed for any time point (blue trace in Figure 5 inset).

**Figure 5.**
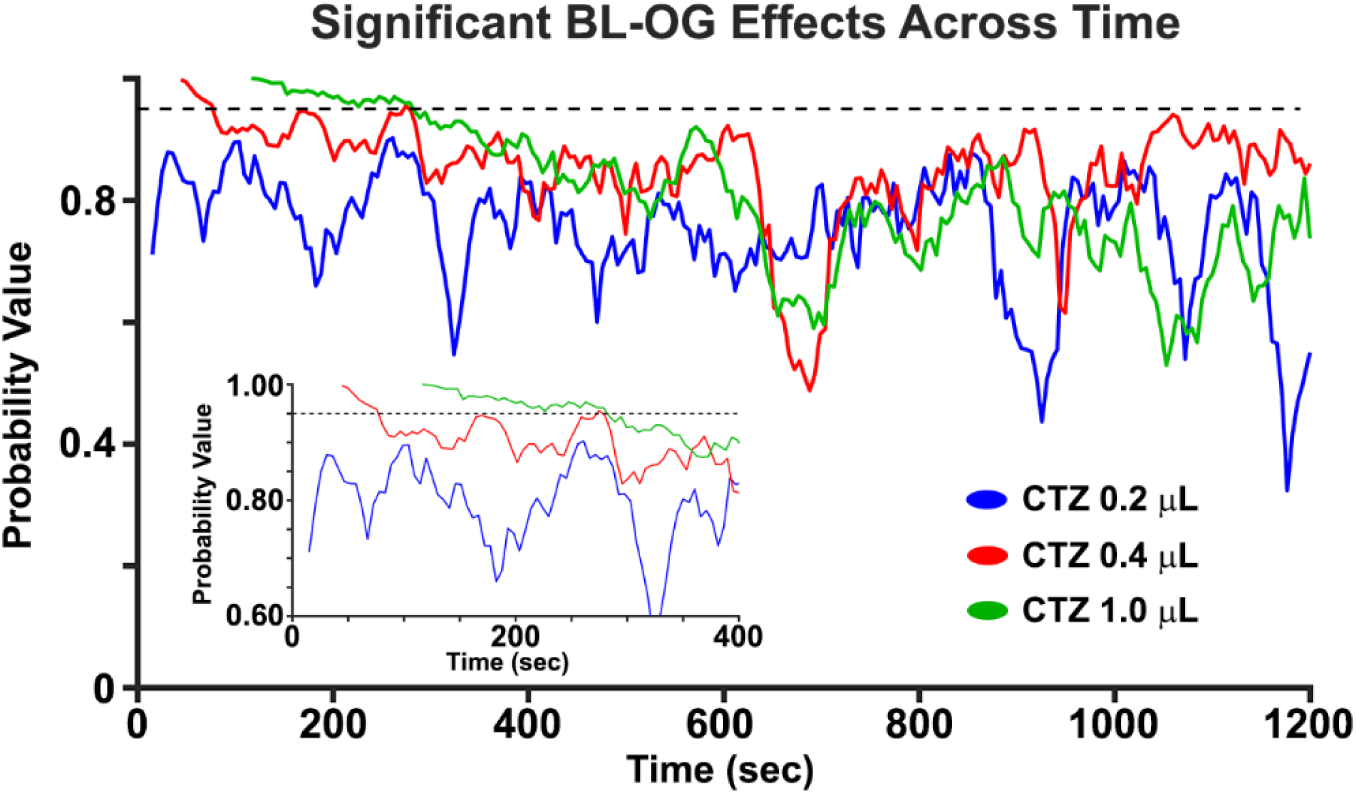
BL-OG effects as a function of time. BL-OG effects on MUA across time for all CTZ doses in the LMO3 group. The inset shows a zoomed in view of the probability values in the first ∼400 secs. Note that the first 24, 48, and 120 seconds are not plotted because of the injection artifact for the 0.2 μL, 0.4 μL, and 1.0 μL, respectively.

## DISCUSSION

We investigated the relationship between bioluminescence and optogenetic excitation of neural activity in anesthetized mice expressing LMO3, and found a systematic relationship between bioluminescence and MUA in animals expressing the LMO3 construct. We also tested the effects of four major elements of the BL-OG reaction on spiking activity (CTZ, CTM, vehicle, and bioluminescence). We failed to observe a dose range where CTM or vehicle modulated MUA. The largest CTZ dose modestly increased neural activity in naïve mice and GLuc cohorts, with no differences in MUA effects between the two groups. These findings suggest that CTZ itself can have non-specific excitatory effects in high concentrations, with selective effects across a large dose range. Taken together, these data indicate that BL-OG is a viable method for regulating activity of genetically-identified neurons *in vivo*. We recommend using a CTZ dose that evokes selective effects, and, when appropriate, testing for possible off target effects when applying this method in new regions and cell types.

### Relationship between bioluminescence and MUA

A major focus of our study was to characterize the relationship between bioluminescence and changes in neural activity relative to CTZ dose. We observed a systematic relationship between bioluminescent activity and MUA (Figure 2B and Figure 3), indicating that the LMO3 molecule robustly drives optogenetic elements *in vivo*. The data also revealed a systematic relationship between bioluminescence and MUA as a function of CTZ dose, with larger infusions of CTZ leading to greater MUA and bioluminescence. The dynamic range of this effect spanned ∼75%, demonstrating that BL-OG effects can be parametrically varied within a large range depending on the CTZ dose used.

Maximal effects of BL-OG were observed during the early phase of CTZ administration (0-400 seconds), and BL-OG effects gradually decreased across time. While reliable bioluminescent signals were observed during 400-800 seconds for all CTZ doses, we failed to find MUA modulations in any CTZ condition at the group level. However, we did observe significant MUA increases in a small subset of animals. Also, for the largest CTZ condition we observed continuous BL-OG effects commencing at the beginning of the CTZ injection and lasting for ∼160 seconds. We found similar, but weaker and shorter, effects for the 0.4 μL CTZ dose, whereas no effects in the 0.2 μL CTZ dose, suggesting that BL-OG effects for these doses require a larger integration time to elicit long-lasting statistically significant modulations.

### Targeted concentration range of CTZ

A major goal of this paper was to assay potential off target effects of four major elements of the BL-OG strategy. Previous chemogenetic studies have shown that driving agents can cause neural artifacts (MacLaren et al., 2016), and light activation can modulate neural activity (Kozai and Vazquez, 2015). We found that neither CTM, the oxidized product of CTZ, nor vehicle modulated neural spiking activity. Similar findings were observed for the lower CTZ doses in both CTZ and GLuc cohorts. Yet, we observed modest increases in MUA in response to the largest CTZ dose.

A key question for all neuromodulation methods is to determine the threshold level at which nonspecific effects emerge. In the case of chemogenetic drivers, the threshold implies a dose at which the activating molecule yields robust responses while producing negligible off target effects. While the specific approach used here (direct neocortical injection) may not be the preferred administration strategy for some BL-OG applications (e.g., in free behavior studies), these results nevertheless indicate a substantial range of CTZ levels across which neocortical nonspecific effects are not a significant concern.

### Advantage of using luciferases to monitor and control neural ensembles

The use of luciferases for studying and manipulating brain circuits has several distinctive advantages over other chemogenetic methods: *(1)* In our BL-OG strategy, a single molecular construct provides both optogenetic and chemogenetic activation options. As such, LMOs can facilitate integrated experiments, whereby a distributed system of neurons can be activated by systemic injection of a chemical, or distinct sub-regions can be controlled by local injections or delivery of light through a fiber optic in the same animal. *(2)* Unlike most chemogenetic options, the rapidly expanding family of optogenetic options in combination with the broad range of existing luciferases and luciferins (Haddock et al., 2010), provide an expanded toolkit to meet specific experimental needs. These options include direct ion channel opening (e.g., using Channelrhodopsin variants) or other light-dependent applications (e.g., engagement of G-protein pathways, as accessed by DREADDs). *(3)* Because distinct luciferins are typically not cross reactive with non-matched luciferases (Haddock et al., 2010), multiple independent BL-OG targets could be independently activated in the same preparation by the choice of luciferins. *(4)* BL-OG has the unique feature of reporting the successful delivery of a pharmacological agent (a luciferin) by emitting light. Even simple optical detection techniques can provide real-time insight into the efficacy and time course of chemical delivered to the preparation (e.g., by peripheral injection). *(5)* Other uses of bioluminescence to study brain functions have been developed (Naumann et al., 2010; Inagaki and Nagai, 2016; Inagaki et al., 2017), providing an even wider range of distinct applications. For instance, bioluminescent molecules have been used as reporters of cell activity in the calcium (Naumann et al., 2010; Saito et al., 2012; Gomez-Ramirez et al., 2017) and voltage domain (Inagaki et al., 2017), and can be tethered to opsins to enable cellular-specific and real-time neural regulation (Pal et al., 2017).

Fiber optic drive of light-sensing molecules can achieve millisecond precision of neural modulation. However, chemogenetic strategies may be more advantageous when sustained modulation of defined populations, especially over widespread regions, is desired. Like most chemogenetic options, BL-OG obviates the need for invasive devices (e.g., implanted fiber optics). Invasive external devices delivering light (e.g., optical fibers) are challenging for many reasons, in part because implants represent a path for pathogens to reach the brain, a concern that is amplified in long-term chronic experiments lasting months or years. In addition, light from optical devices can be limited in its utility for continuously exciting (or inhibiting) a population for extended periods of time because power from photon emission can overheat tissue and lead to photodamage and/or photobleaching (Denk and Svoboda, 1997; Chirico et al., 2003; Pashaie and Falk, 2013). Also, optical implants can be punitive when targeting disparate foci because insertion of several fibers can cause considerable tissue damage that lead to behavioral deficits. Chemogenetic solutions to this latter drawback are particularly encouraging because the ability to concomitantly modulate large-scale, but functionally connected, cell ensembles is fundamental for interrogating neural codes underlying complex behavior of mammalian networks.

## Supplementary material

None

## ACKNOWLEDGEMENTS

We would like to thank Dr. Ken Berglund for providing the plasmid for the NonActive LMO3 (R115A) construct.

## REFERENCES

Berglund K, Birkner E, Augustine GJ, Hochgeschwender U (2013) Light-emitting channelrhodopsins for combined optogenetic and chemical-genetic control of neurons. PLoS One 8:e59759.

Berglund K, Fernandez AM, Gutekunst CN, Hochgeschwender U, Gross RE (2019) Step-function luminopsins for bimodal prolonged neuromodulation. J Neurosci Res.

Berglund K, Tung JK, Higashikubo B, Gross RE, Moore CI, Hochgeschwender U (2016a) Combined Optogenetic and Chemogenetic Control of Neurons. Methods Mol Biol 1408:207–225.

Berglund K, Clissold K, Li HE, Wen L, Park SY, Gleixner J, Klein ME, Lu D, Barter JW, Rossi MA, Augustine GJ, Yin HH, Hochgeschwender U (2016b) Luminopsins integrate opto- and chemogenetics by using physical and biological light sources for opsin activation. Proc Natl Acad Sci U S A 113:E358–367.

Birkner E, Berglund K, Klein ME, Augustine GJ, Hochgeschwender U (2014) Non-invasive activation of optogenetic actuators. Proc SPIE Int Soc Opt Eng 8928.

Boyden ES, Zhang F, Bamberg E, Nagel G, Deisseroth K (2005) Millisecond-timescale, genetically targeted optical control of neural activity. Nat Neurosci 8:1263–1268.

Cardin JA, Carlen M, Meletis K, Knoblich U, Zhang F, Deisseroth K, Tsai LH, Moore CI (2009) Driving fast-spiking cells induces gamma rhythm and controls sensory responses. Nature 459:663–667.

Chirico G, Cannone F, Baldini G, Diaspro A (2003) Two-photon thermal bleaching of single fluorescent molecules. Biophys J 84:588–598.

Chou WC, Liao KW, Lo YC, Jiang SY, Yeh MY, Roffler SR (1999) Expression of chimeric monomer and dimer proteins on the plasma membrane of mammalian cells. Biotechnol Bioeng 65:160–169.

Denk W, Svoboda K (1997) Photon upmanship: why multiphoton imaging is more than a gimmick. Neuron 18:351–357.

Fenno L, Yizhar O, Deisseroth K (2011) The development and application of optogenetics. Annu Rev Neurosci 34:389–412.

Gomez-Ramirez M, Trzcinski NK, Mihalas S, Niebur E, Hsiao SS (2014) Temporal correlation mechanisms and their role in feature selection: a single-unit study in primate somatosensory cortex. PLoS Biol 12:e1002004.

Gomez-Ramirez M, More IA, Pal A, Connors BW, Kauer JA, Lipscombe D, Hochgeschwender U, Moore CI (2017) Imaging and regulation of cortical neurons using bioluminescent molecules: A biological method for tracking neural dynamics and driving optogenetic elements in vivo. Soc Neurosci Abstr.

Gomez JL, Bonaventura J, Lesniak W, Mathews WB, Sysa-Shah P, Rodriguez LA, Ellis RJ, Richie CT, Harvey BK, Dannals RF, Pomper MG, Bonci A, Michaelides M (2017) Chemogenetics revealed: DREADD occupancy and activation via converted clozapine. Science 357:503–507.

Haddock SH, Moline MA, Case JF (2010) Bioluminescence in the sea. Ann Rev Mar Sci 2:443–493.

Inagaki S, Nagai T (2016) Current progress in genetically encoded voltage indicators for neural activity recording. Curr Opin Chem Biol 33:95–100.

Inagaki S, Tsutsui H, Suzuki K, Agetsuma M, Arai Y, Jinno Y, Bai G, Daniels MJ, Okamura Y, Matsuda T, Nagai T (2017) Genetically encoded bioluminescent voltage indicator for multi-purpose use in wide range of bioimaging. Sci Rep 7:42398.

Kato HE, Zhang F, Yizhar O, Ramakrishnan C, Nishizawa T, Hirata K, Ito J, Aita Y, Tsukazaki T, Hayashi S, Hegemann P, Maturana AD, Ishitani R, Deisseroth K, Nureki O (2012) Crystal structure of the channelrhodopsin light-gated cation channel. Nature 482:369–374.

Kim CK, Adhikari A, Deisseroth K (2017) Integration of optogenetics with complementary methodologies in systems neuroscience. Nat Rev Neurosci 18:222–235.

Knopfel T, Lin MZ, Levskaya A, Tian L, Lin JY, Boyden ES (2010) Toward the second generation of optogenetic tools. J Neurosci 30:14998–15004.

Kozai TD, Vazquez AL (2015) Photoelectric artefact from optogenetics and imaging on microelectrodes and bioelectronics: New Challenges and Opportunities. J Mater Chem B Mater Biol Med 3:4965–4978.

MacLaren DA, Browne RW, Shaw JK, Krishnan Radhakrishnan S, Khare P, Espana RA, Clark SD (2016) Clozapine N-Oxide Administration Produces Behavioral Effects in Long-Evans Rats: Implications for Designing DREADD Experiments. eNeuro 3.

Naumann EA, Kampff AR, Prober DA, Schier AF, Engert F (2010) Monitoring neural activity with bioluminescence during natural behavior. Nat Neurosci 13:513–520.

Nichols CD, Roth BL (2009) Engineered G-protein Coupled Receptors are Powerful Tools to Investigate Biological Processes and Behaviors. Front Mol Neurosci 2:16.

Pal A, Gomez-Ramirez M, Medendrop WE, Zaidi Z, Kauer JA, Lipscombe D, Connors BW, Moore CI, Hochgeschwender U (2017) Characterization, sub-cellular targeting and novel applications of a split Gaussia luciferase based genetically encoded calcium indicator. Soc Neurosci Abstr.

Park SY, Song SH, Palmateer B, Pal A, Petersen ED, Shall GP, Welchko RM, Ibata K, Miyawaki A, Augustine GJ, Hochgeschwender U (2017) Novel luciferase-opsin combinations for improved luminopsins. J Neurosci Res.

Pashaie R, Falk R (2013) Single optical fiber probe for fluorescence detection and optogenetic stimulation. IEEE Trans Biomed Eng 60:268–280.

Prakash M, Medendorp WE, Hochgeschwender U (2018) Defining parameters of specificity for bioluminescent optogenetic activation of neurons using in vitro multi electrode arrays (MEA). J Neurosci Res.

Roth BL (2016) DREADDs for Neuroscientists. Neuron 89:683–694.

Saito K, Chang YF, Horikawa K, Hatsugai N, Higuchi Y, Hashida M, Yoshida Y, Matsuda T, Arai Y, Nagai T (2012) Luminescent proteins for high-speed single-cell and whole-body imaging. Nat Commun 3:1262.

Tung JK, Gutekunst CA, Gross RE (2015) Inhibitory luminopsins: genetically-encoded bioluminescent opsins for versatile, scalable, and hardware-independent optogenetic inhibition. Sci Rep 5:14366.

Tung JK, Shiu FH, Ding K, Gross RE (2018) Chemically activated luminopsins allow optogenetic inhibition of distributed nodes in an epileptic network for non-invasive and multi-site suppression of seizure activity. Neurobiol Dis 109:1–10.

Urban DJ, Roth BL (2015) DREADDs (designer receptors exclusively activated by designer drugs): chemogenetic tools with therapeutic utility. Annu Rev Pharmacol Toxicol 55:399–417.

Welsh JP, Patel KG, Manthiram K, Swartz JR (2009) Multiply mutated Gaussia luciferases provide prolonged and intense bioluminescence. Biochem Biophys Res Commun 389:563–568.

Zenchak JR, Palmateer B, Dorka N, Brown TM, Wagner LM, Medendorp WE, Petersen ED, Prakash M, Hochgeschwender U (2018) Bioluminescence-driven optogenetic activation of transplanted neural precursor cells improves motor deficits in a Parkinson’s disease mouse model. J Neurosci Res.

Zhang F, Wang LP, Boyden ES, Deisseroth K (2006) Channelrhodopsin-2 and optical control of excitable cells. Nat Methods 3:785–792.

Zhang F, Prigge M, Beyriere F, Tsunoda SP, Mattis J, Yizhar O, Hegemann P, Deisseroth K (2008) Red-shifted optogenetic excitation: a tool for fast neural control derived from Volvox carteri. Nat Neurosci 11:631–633.

Zhu Y, Feng B, Schwartz ES, Gebhart GF, Prescott SA (2015) Novel method to assess axonal excitability using channelrhodopsin-based photoactivation. J Neurophysiol 113:2242–2249.

